# Stratification of the Gut Microbiota Composition Landscape Across the Alzheimer’s Disease Continuum in a Turkish Cohort

**DOI:** 10.1101/2021.10.28.466378

**Authors:** Süleyman Yıldırım, Özkan Ufuk Nalbantoğlu, Abdulahad Bayraktar, Fatma Betül Ercan, Aycan Gündoğdu, Halil Aziz Velioğlu, Mehmet Fatih Göl, Ayten Ekinci Soylu, Fatma Koç, Ezgi Aslan Gürpınar, Kübra Sogukkanlı Kadak, Muzaffer Arıkan, Adil Mardinoğlu, Mehmet Koçak, Emel Köseoğlu, Lütfü Hanoğlu

## Abstract

Alzheimer’s disease (AD) is a heterogeneous neurodegenerative disorder that spans over a continuum with multiple phases including preclinical, mild cognitive impairment, and dementia. Unlike most other chronic diseases there are limited number of human studies reporting on AD gut microbiota in the literature. These published studies suggest that the gut microbiota of AD continuum patients varies considerably throughout the disease stages, raising expectations for existence of multiple microbiota community types. However, the community types of AD gut microbiota were not systematically investigated before, leaving important research gap for diet-based intervention studies and recently initiated precision nutrition approaches aiming at stratifying patients into distinct dietary subgroups. Here, we comprehensively assessed the community types of gut microbiota across the AD continuum. We analyze 16S rRNA amplicon sequencing of stool samples from 27 mild cognitive patients, 47 AD, and 51 non-demented control subjects using tools compatible with compositional nature of microbiota. To characterize gut microbiota community types, we applied multiple machine learning techniques including partitioning around the medoid clustering, fitting probabilistic Dirichlet mixture model, Latent Dirichlet Allocation model, and performed topological data analysis for population scale microbiome stratification based on Mapper algorithm. These four distinct techniques all converge on Prevotella and Bacteroides partitioning of the gut microbiota across AD continuum while some methods provided fine scale resolution in partitioning the community landscape. The Signature taxa and neuropsychometric parameters together robustly classify the heterogenous groups within the cohort. Our results provide a framework for precision nutrition approaches and diet-based intervention studies targeting AD cohorts.

**IMPORTANCE:** The prevalence of AD worldwide is estimated to reach 131 million by 2050. Most disease modifying treatments and drug trials have failed due partly to the heterogeneous and complex nature of the disease. Unlike other neurodegenerative diseases gut microbiota of AD patients is poorly studied. Recently initiated ambitious precision nutrition initiative or other diet-based interventions can potentially be more effective if the heterogeneous disease such as AD is deconstructed into multiple strata allowing for better identification of biomarkers across narrower patient population for improved results. Because gut microbiota is inherently integral part of the nutritional interventions there is unmet need for microbiota-informed stratification of AD clinical cohorts in nutritional studies. Our study fills in this gap and draws attention to the need for microbiota stratification as one of the essential steps for precision nutrition interventions. We demonstrate that while Prevotella and Bacteroides clusters are the consensus partitions the newly developed probabilistic methods can provide fine scale resolution in partitioning the AD gut microbiome landscape.

## INTRODUCTION

Alzheimer’s Disease (AD) is the most common form of dementia worldwide and its prevalence is estimated to reach 131 million by 2050 [1]. AD spans over a continuum starting with the non-symptomatic pre-clinical stage and advancing through the spectrum of clinical stages. These stages are dashed with distinct pathophysiological states [2], namely the amyloid-tau-neuroinflammation axis. The clinical continuum entails mild memory loss and/or cognitive impairments (mild cognitive impairment, MCI due to AD) and trajectories for function leading to memory problems besides cognitive impairment (dementia phase); and finally complete loss of independent functioning towards the end stage [3]. Moreover, The Alzheimer’s dementia phase is further broken down into the stages of mild, moderate and severe, thereby making AD a complex and highly heterogenous disease.

Traditionally, pathogenesis of AD is attributed to extracellular aggregation of amyloid-β-peptides (Aβ) in senile plaques and intracellular depositions of hyperphosphorylated tau that forms neurofibrillary tangles [4]. Although numerous clinical trials based on the amyloid postulates have been attempted virtually all of them have failed [5]. The unsettlingly consistent failure of clinical trials targeting single target amyloid pathways prompted researchers to refine the amyloid hypothesis [6] and even extend it to periphery [7]. Recently, a group of AD researchers asserted that infectious agents reach and remain dormant in the central nervous system (CNS) and undergo reactivation during aging, sparking cascades of inflammation, induce Aβ, and ultimately neuronal degeneration [8]. Chronic inflammation in CNS mediated by microglial toxicity as well as systemic inflammation in the periphery is widely recognized in AD and linked to amyloid cascade hypothesis in animal experiments [9, 10]. None of the drugs available today for Alzheimer’s dementia slow or stop the damage and destruction of neurons [11]. Intervention at different points along the Alzheimer’s continuum should therefore be multimodal and involve targeting neuropathology in brain, systemic inflammation in the body, and metabolic processes in the periphery that escalate the disease in brain [12]. Non-pharmacologic, targeted, personalized, and multimodal disease modifying interventions in AD, including diet and lifestyle changes to optimize metabolic parameters has recently been under investigation [13–16].

A growing body of evidence suggest that human gut microbiota is strongly associated with human metabolic processes in all organs including brain [17] and implicated in neuroinflammation via brain-gut axis [18]. Gut microbes across animal models influence CNS by modulation of neuroimmune function, sensory neuronal signaling, and metabolic activity [19]. Several studies using transgenic animal model of AD reported gut microbiota alterations (see [19]) but these animal models poorly mirror human AD. Unexpectedly, only a few human clinical studies on AD were reported in the literature [20–28]. Of these studies, gut microbiota associated metabolites such elevated Trimethylamine N-oxide (TMAO) in CSF [26] and altered bile acids profile [28] were directly implicated in AD dementia. Importantly, dietary pattern of AD patients is at the center of the precision medicine approaches [29]. Also, diet is one of the most important factors modulating gut microbiota-based active metabolites. Disease modifying approaches involving diet should therefore consider microbiota in AD. Indeed, a recent study [23] tested the impact of a modified Mediterranean ketogenic diet on gut microbiome composition and demonstrated that the diet can modulate the gut microbiome and metabolites in association with improved AD biomarkers in CSF. These published studies, however, did not comprehensively investigate AD microbiota subclusters across the disease continuum, leaving important gap in our understanding of human microbiota in a highly heterogenous disease. Recently initiated ambitious precision nutrition approaches [30–33] cannot be applied on a highly heterogenous disease before deconstructing the disease into multiple strata and tailoring therapies accordingly.

In the present study, we postulated that gut microbiota dysbiosis along the AD continuum should reflect an overlapping yet distinct community types. We show that AD gut microbiota includes distinct community types and the cognitive impairments in AD continuum is associated with unique gut microbiota signatures. Elucidating the diversity and community types of gut microbiota would facilitate identification of stratification biomarkers thereby contributing to precision nutrition approaches in AD.

## RESULTS

### Study Design and Participant Characteristics

The study cohort consisted of 47 AD, 27 MCI (all amnestic), and 51 subjects non-demented controls (N=125). To minimize dietary confounding effect on the microbiome, we included healthy co-habiting spouses of the patients sharing the same diet as controls. The control group therefore largely (n=27) comprised partners of the patients. Participants were recruited in two health centers located in different cities. The cohort groups were statistically not different in terms of sex, but age and education factors were significantly different (Table 1), therefore statistically adjusted in analyses. Expectedly, the groups were also different in cognitive tests including the Mini-Mental State Exam (MMSE), and the Clinical Dementia rating (CDR). Most AD participants had very mild or mild dementia, with clinical dementia rating (CDR) scores ranging from 0.5–3 (median CDR 1 for AD; 0.5 for MCI and 0 for the control group). The median MMSE scores were significantly higher in control (MMSE=27) and MCI (MMSE=26) groups than AD (MMSE=16). A subset of AD patients (n=12) was clinically asked to undergo lumbar puncture to ascertain diagnosis using CSF biomarkers including Aβ42/Aβ40 ratio, phosphorylated tau (p-tau), and the p-tau/Aβ42 ratio (Supplementary Table S1). We collected medication information from the patient’s registry.

**Table 1.**
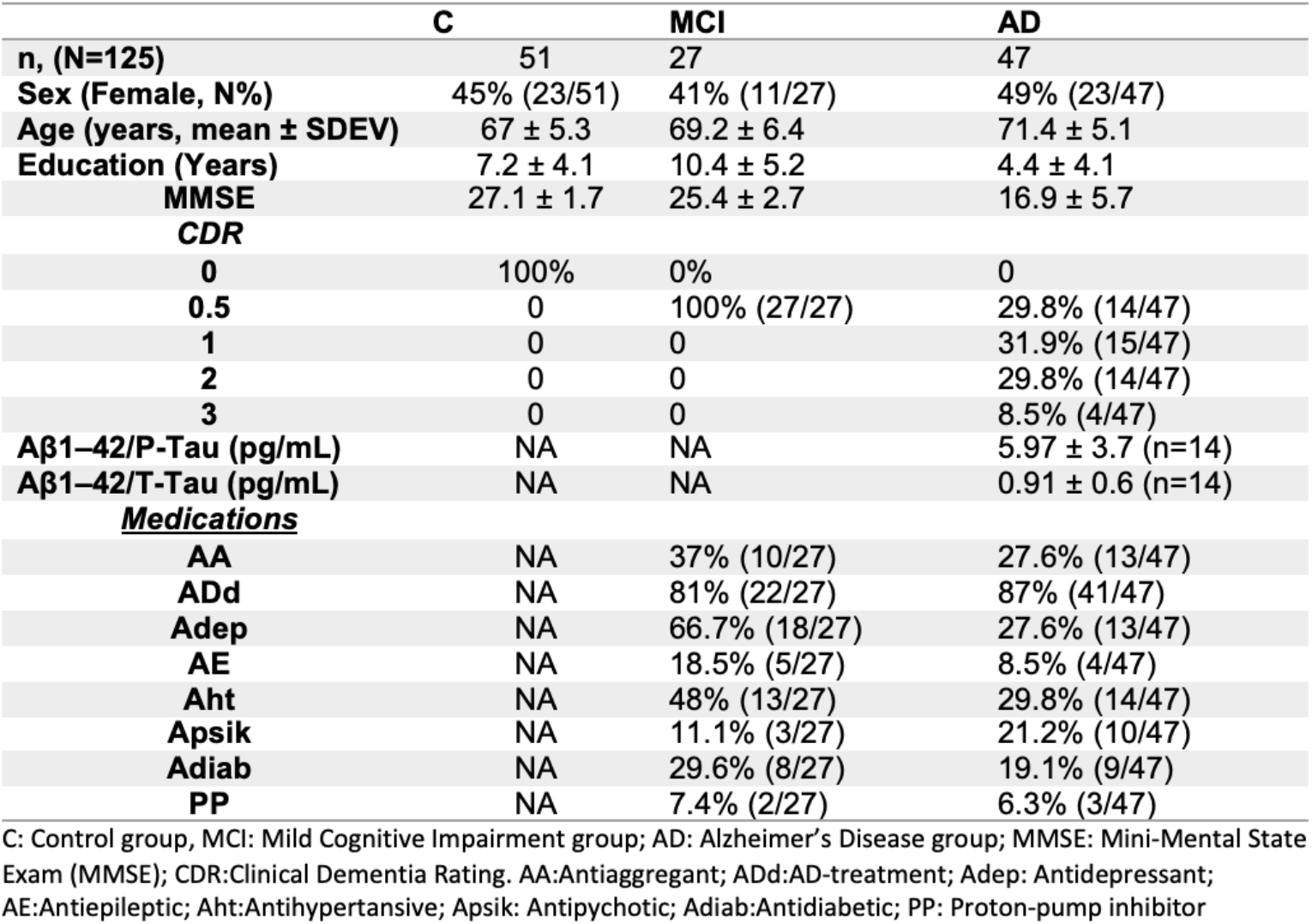
Demographic characteristics of the participants in the cohort.

### Microbiome composition is associated with disease status along the AD continuum

The gut microbiota was profiled using the V3-V4 hypervariable region of the 16S rRNA gene; The Nephele automatic pipeline denoised the paired-end sequences and assigned amplicon sequence variants (ASVs) according to DADA2 [34]. The Nephele produced both unrarefied and the rarefied ASV tables. The rarefied table included a total of 3486 ASVs in the table (10769 sequences/sample) for downstream analyses.

The phylum level taxonomic analysis showed typical human gut microbiota profile in terms of over-abundance of *Firmicutes, Bacteroidetes, and Proteobacteria* (Figure 1a). Together with *Verrucomicrobia*, and *Actinobacteria* the five phyla comprised 99% of all reads but *Proteobacteria* was overrepresented in AD patient samples. Notably, the genus level relative abundance distributions across samples showed *Prevotella_9* and *Bacteroides* were the most abundant of top30 genera across the samples (Figure 1b). To perform differential abundance analysis between samples we sought concordance analysis among multiple tools. ANCOM-BC or ALDEx2, when used covariates in their models, both agreed that only *Ruminoccus_unclassified* is significantly differentially abundant among the groups (data not shown). Nevertheless, when we employed limma-voom R package (age and sex adjusted, FDR<0.05) we found that *Prevotella_9, Bacteroides* and members of *Ruminococcaceae* family were among the top most significant differentially abundant taxa (ASV) between the cohort groups (Supplementary Tables S2-5). A comprehensive comparative statistical assessment of multivariate and compositional methods [35] demonstrated ALDEx2 or alike tools suffer from low power while limma-voom and songbird in their own class were the best performers.

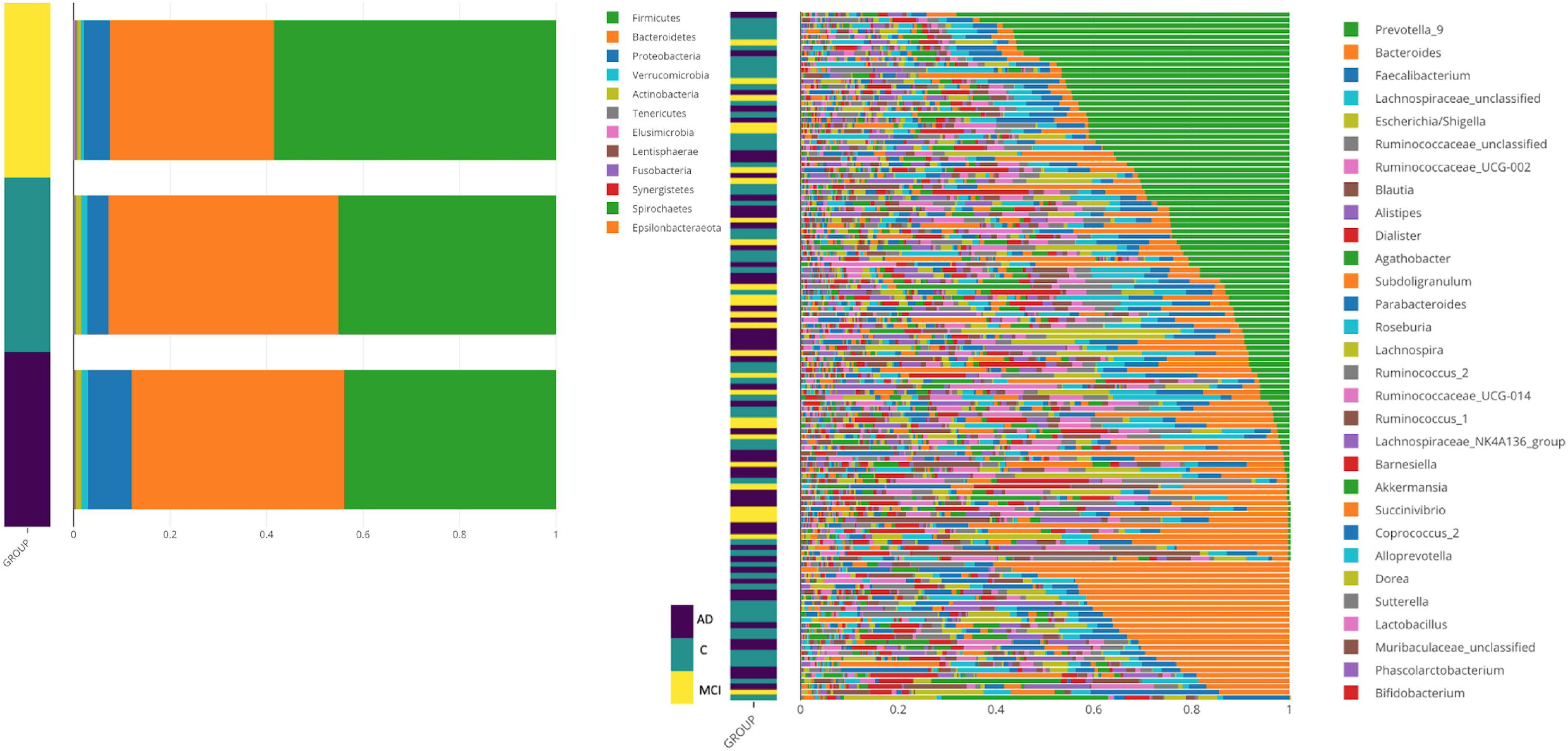

Alpha diversity indices (Shannon, Inverse Simpson) did not show significant differences after multiple testing corrections (Kruskal-Wallis, Supplementary Figure S1 (a-d), FDR>0.05) but richness index, Chao1, showed significant difference between MCI and the control group (pairwise Wilcoxon rank sum test, p=0.008074).

We employed both relative abundances based and recently developed compositionally aware tools, namely DEICODE [36] and Songbird [37] to compare the composition and structure of bacterial communities in samples using multiple beta diversity indices (Bray-Curtis, Jaccard, and Aitchison). The principal coordinates analysis showed separation of the three groups by both Bray-Curtis and Jaccard indices (Figure 2a-b). We used adonis2 function in qiime2 plugin (q2-diversity) to perform PERMONAVA analysis with 999 permutations and included interaction terms (Supplementary Table S6) and seperation of the groups were highly significant (P=0.0001). Age and Sex also significantly contributed to the total variance (P<0.001) but the interaction terms were not significant. Furthermore, dispersion between groups tests (PERMDISP) indicated only the dispersion MCI group is significantly heteregenous (pairwise comparisons p=0.033 for AD-MCI; p=0.024 for C-MCI; p=0.672 for AD-C), which may be attributed to unbalanced design. We added further support for the seperation of the three groups from other ordinations. The Canonical Analysis of Principal Coordinates (CAP) analysis unambigiously showed the three groups are distinct (Figure 2c, trace statistic = 0.86855, p=0.001, 999 permutations). The final support in beta diversity was provided by the DEICODE analysis (robust Aitchison PCA) (Figure 2d, PERMANOVA p=0.02), which indicated that the three groups are distinct, and the community clusters are largely driven by a subset of ASVs with taxonomic assignment *Prevotella_9, Bacteroides*, a unclassified genus within *Ruminococcaceae* family (*Ruminococcaceae_unclassified*), and *Escherichia/Shigella*. Moreover, the co-occurrence analysis using SparCC showed that *Prevotella_9* and *Bacteroides* were negatively correlated (Correlation=-0.4445, FDR =0.09355). Moreover, the genus level PCoAs showed partially overlapping clusters of these two taxa while the groups overall were also significantly separated (PERMANOVA, p <0.0001, Supplementary Figures S2 (a-c)). We therefore placed particular attention to these two taxa in the rest of the downstream analyses.

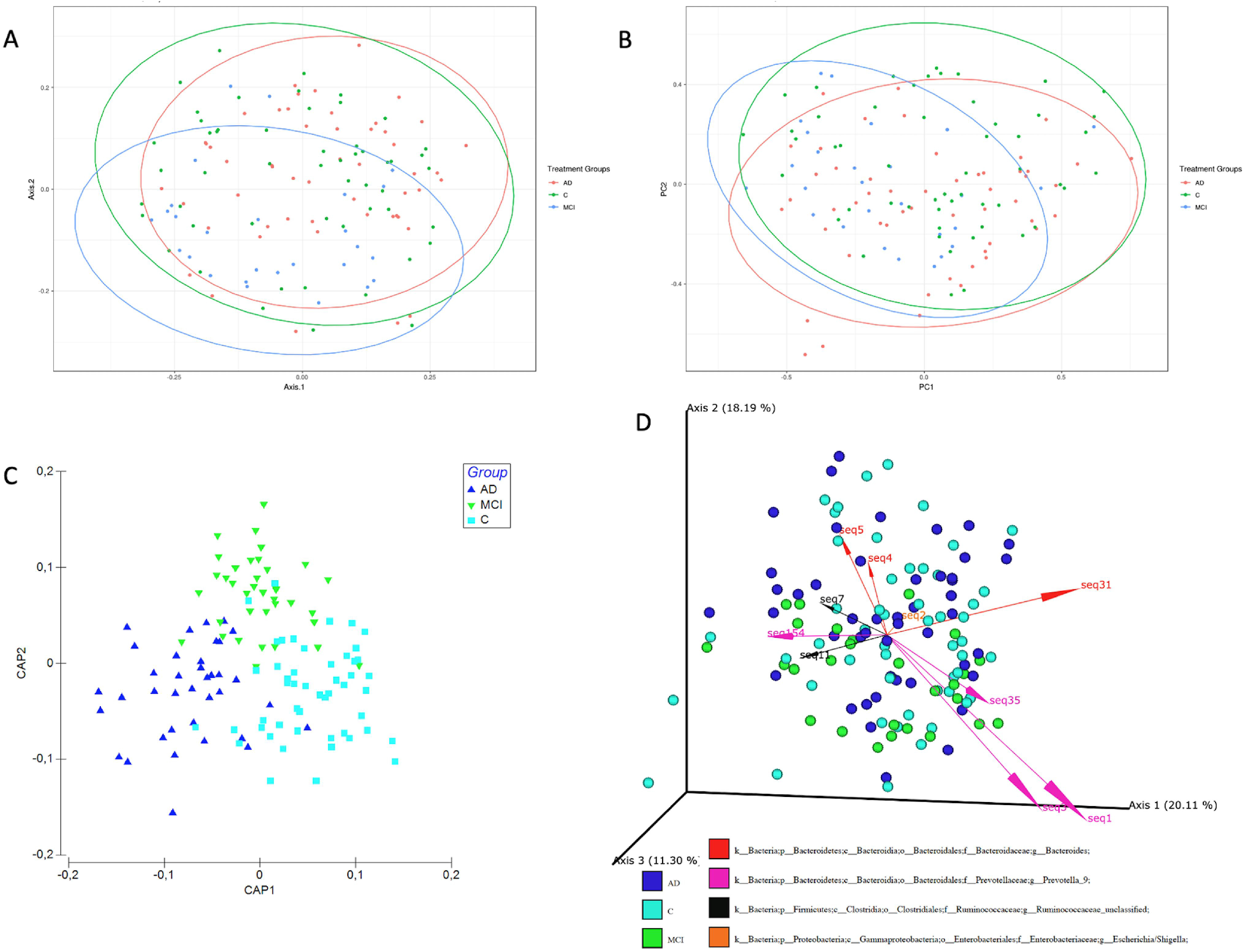

Enrichment analysis by multinomial regression embedded in the songbird tool with regard to covariates (formula: Age+Sex+Edu+MMSE+CDR+Groups(levels=(“C”, “MCI”,”AD”)) indicated that the natural log ratio of *Prevotella_9* to *Bacteroides* and *Prevotella_9* to *Escherichia/Shigella* significantly separated AD group from the control group (Welch’s t-test, FDR adjusted p=0.04) but not from the MCI group (Figure 3a-d). Importantly, the songbird excluded 25 samples from this analysis due to zero-rich abundances that do not allow for center-log ratio calculations. We therefore tested the natural log ratio of top 25% allowing to include all samples in the analysis (“Set1” in Supplementary Table S7) to the bottom 25% (“Set2”, Supplementary Table S8) of the ranked ASVs associated with the AD relative to the control group; also, same ratios for MCI relative to the control group (“Set3” and “Set4”, Supplementary Table S8) and the ASVs enriched in each group were visualized with Qurro [38]. Both sets of ranked log ratios revealed significant differences (Graph Pad Prism) between the log ratios of features differentiating groups (Welch’s t-test, FDR adjusted p= 0.0002).

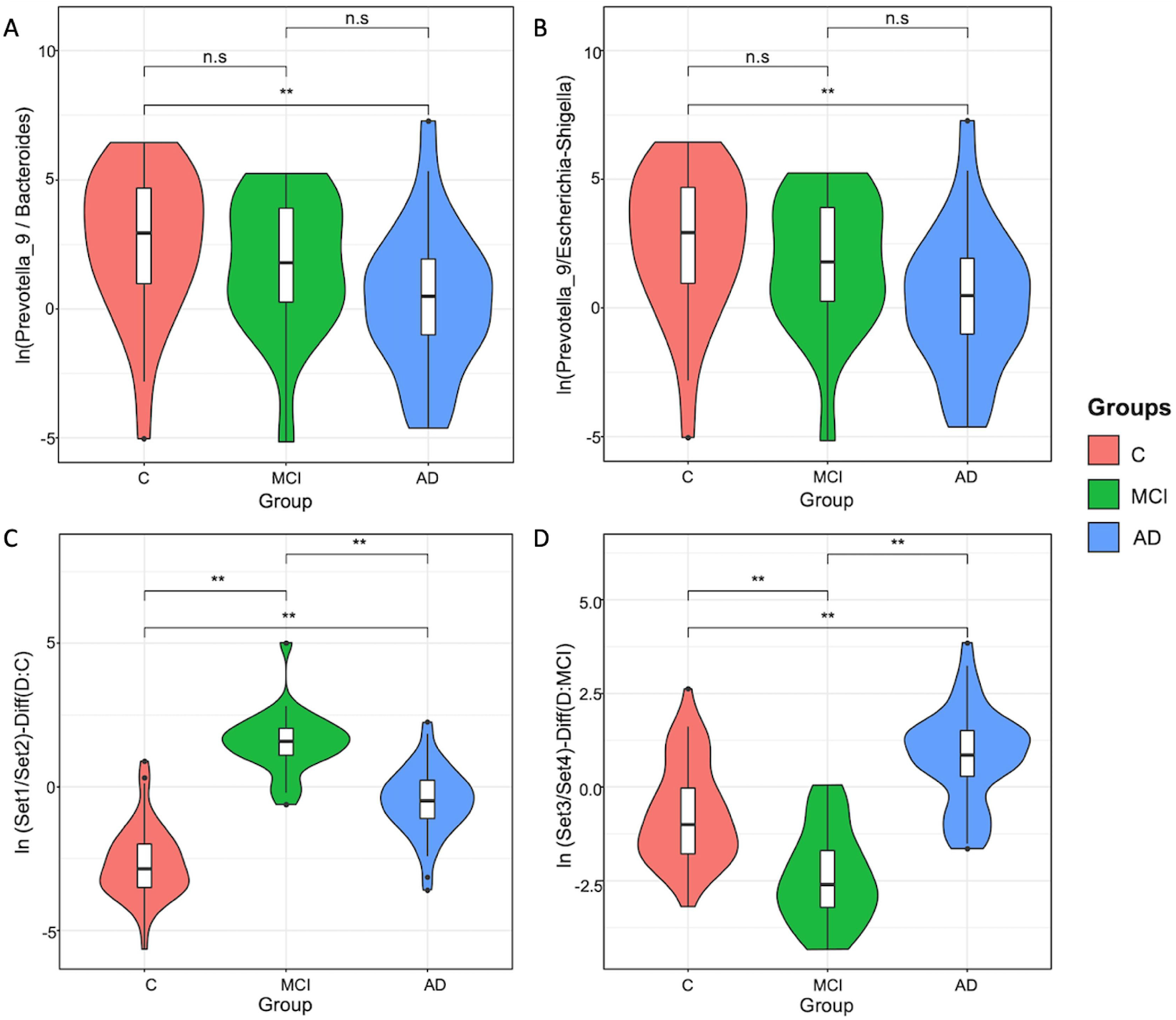

### Discrete multiple subsets of gut microbiota exist along the AD continuum

Considering the preceding results, we postulated that gut microbiota profile along the AD continuum does not represent a single state, rather, distinct yet overlapping community types. We addressed this hypothesis using four unique methods: 1-Partitioning around medoid (PAM)-based clustering [39], 2-Fitting Dirichlet multinomial mixture (DMM) models to partition microbial community profiles into a finite number of clusters [40] using the Laplace approximation, 3-Fitting Latent Dirichlet Allocation (LDA) [41, 42] using perplexity measure, and 4-Analyzing topological futures of data density [43] based on the *Mapper* algorithm to capture subtle and non-linear patterns of high-dimensional datasets and population level stratification.

The PAM-based clustering identified three (k=3) distinct clusters based on Gap statistics (Supplementary Figure S3a). PCoA analysis of the sample abundances in the three clusters indicated significant separation of the clusters (Figure 4a, PERMANOVA, p=0.001). We confirmed optimum number of clusters using both Jensen-Shannon and Bray-Curtis distance metrices (data not shown). The relative abundance of the genus *Prevotella_9* dominated cluster-1 while the genus *Bacteroides* showed the highest relative abundances in the other two clusters (Figure 4b).

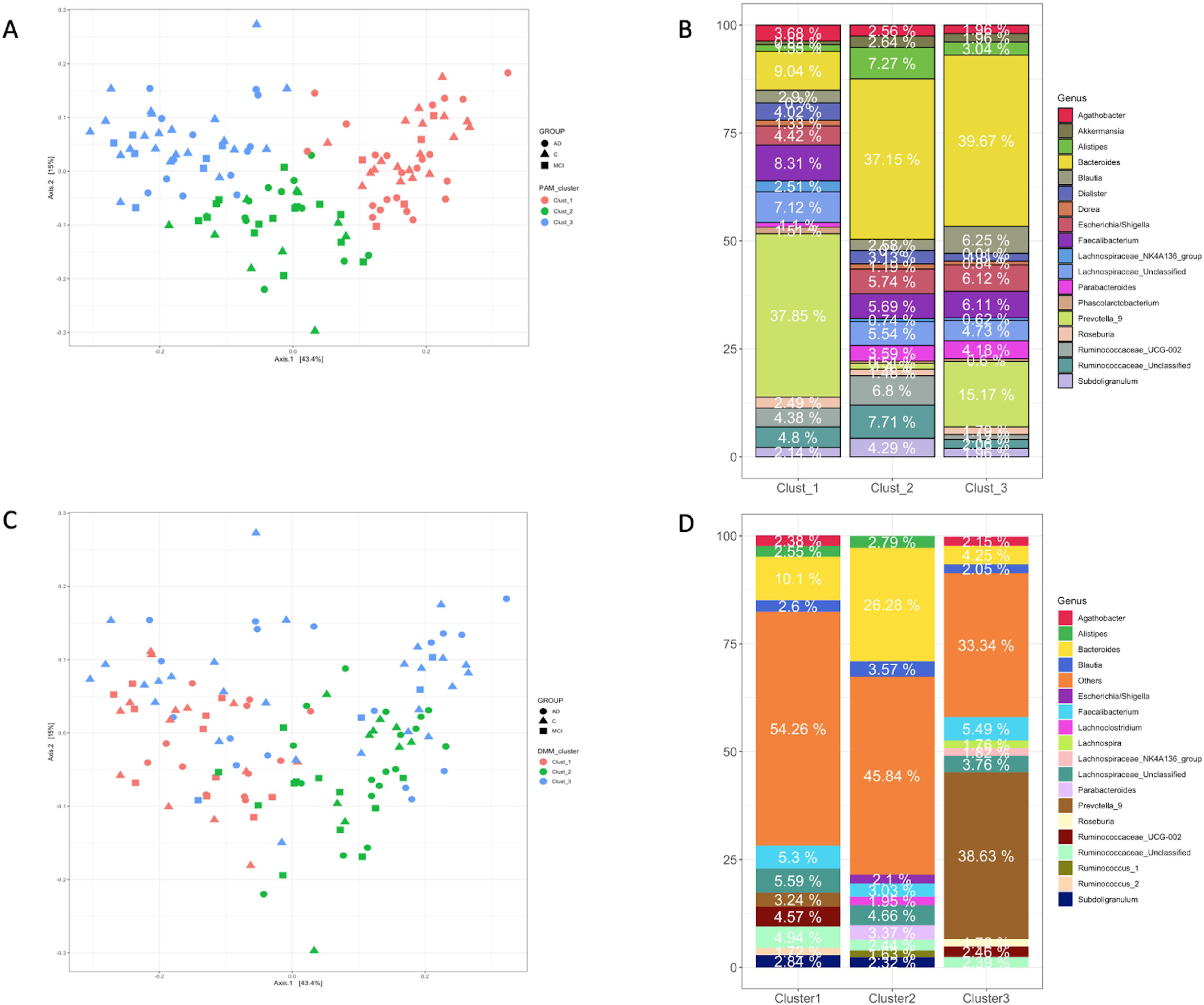

Next, we employed the Dirichlet multinomial mixtures probabilistic community modeling using the *DirichletMultinomial* R package [40] and fitting genus level absolute abundances. Based on Laplace approximation three clusters (cluster 1, 2, and 3) represented the best model fit (Supplementary Figure 3b), which was congruent with the PAM-based clustering. The PCoA analysis of these clusters and PERMANOVA pairwise tests further supported existence of three distinct clusters within the microbial community (Figure 4c, PERMANOVA, p=0.01). The genus *Bacteroides* was the most abundant taxa in the first two clusters and the third cluster was dominated by *Prevotella_9* (Figure 4d). Notably, cluster2 included significantly higher abundance of *Bacteroides* (26.3%) than cluster1 (9.9%) and cluster3 (4.7%). In addition to highly enriched *Bacteroides* in cluster2 the decreasing trend of *Faecalibacterium* abundance and elevated abundance of inflammation associated *Escherchia/Shigella* suggested that cluster2 can be named *“Bacteroides2 (Bact2) enterotype”* as recently described [44, 45]. Reportedly, abundance of *Bacteroides* in Bact2 enterotype can reach as high as 78% in patients with inflammatory bowel disease and is associated with systemic inflammation. These results suggest that cluster2 includes patients with aggravated systemic inflammation.

We also performed SIMPER analysis based on Bray-Curtis distance to identify taxa contributing most to dissimilarities between clusters (data not shown). *Bacteroides, Prevotella_9, Faecalibacterium*, and taxa within *Ruminococcaceae* family ranked among the top ten taxa contributing most to differences between the three DMM clusters. To examine which factors were associated with the DMM clusters we analyzed distribution of clinical metadata and diversity metrics within the clusters. Alpha diversity indices (Chao1, Shannon, and Inverse Simpson) were statistically different between all three clusters after Benjamini-Hochberg FDR adjustment. However, CDR, MMSE, Age, Sex, and Education were not significant between the clusters (Kruskal Wallis test followed by Dunn’s posthoc test, FDR<0.05 and Fisher’s Exact test was used for Sex parameter). (Figure 5 a-h)

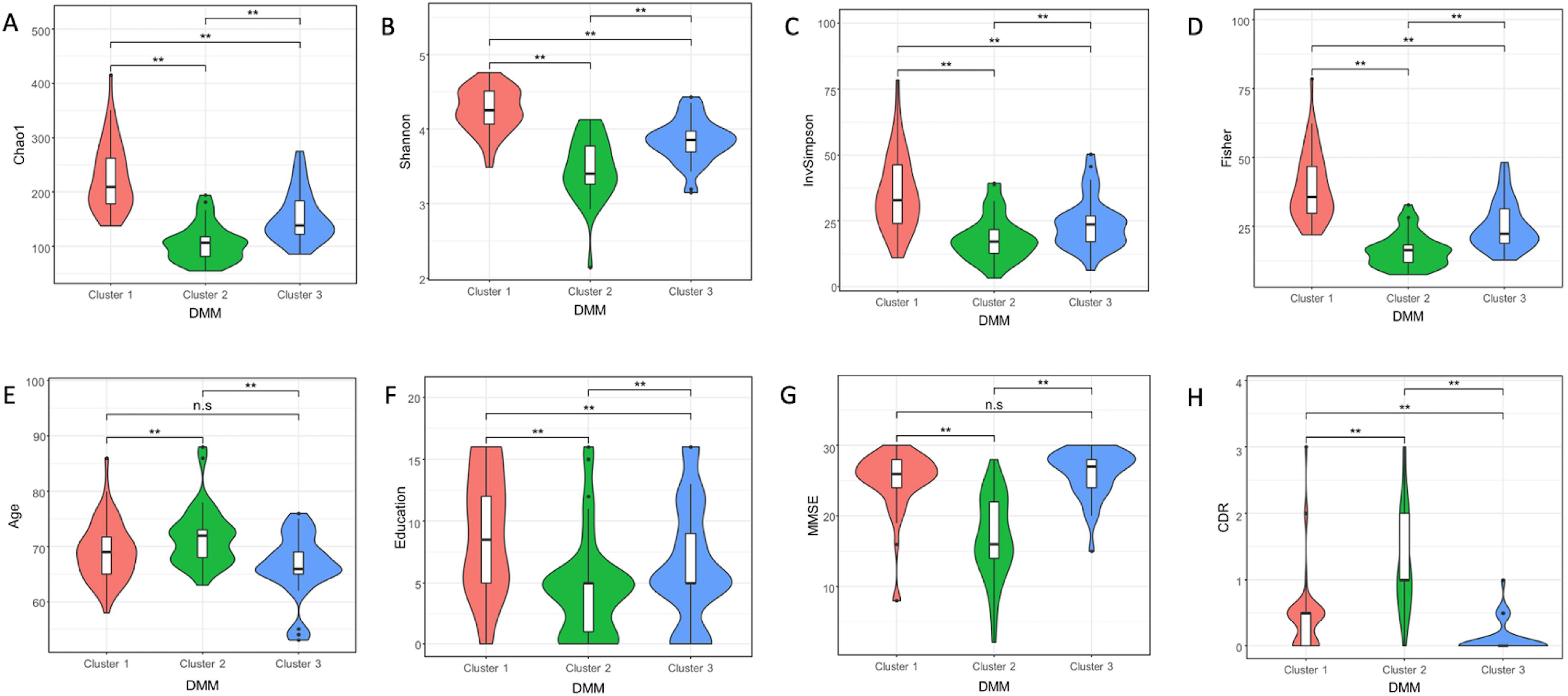

We next tested LDA potential to stratify gut microbiota of the cohort participants. This unsupervised machine learning technique is increasingly finding acceptance in the field of microbiome [46–48] for its unique ability to reveal latent or hidden groups within the data cloud. Supplementary Figure S4 shows LDA model’s perplexity parameter and log-likelihood values to find optimal number of clusters. Both parameters continued to partition the community without reaching a clear optimum. This finding is unexpectedly consistent with recent publications using LDA in microbial ecology [46–48]. Bacteria probability distributions (ranked by probability ≥ 1% in descending order) across the subgroups are displayed in Figure 6a. Interstingly, of the ten subgroups two subgroups were dominated by *Bacteroides* (topic1 and topic5) and a subgroup (topic2) dominated by *Prevotella_9* with 97% probability. These subgroups therefore resemble subgroups detected by PAM and DMM in terms of prevalence of *Bacteroides* and *Prevotella_9*. Unlike DMM and PAM, however, LDA detected a distinct subgroup (topic10) with top ranking genus was *Escherichia/Shigella*, which also included putatively opportunistic bacteria such as *Entercoccus* and *Klebsiella*. Subgroups 4, 6, and 9 were conspicious with the genera known to produce butyrate and acetate or is mucinphilic. Even though we present first ten subgroups (topics) here we also examined higher order subgroups and observe that the ten subgroups are further partitioned into additional subgroups such as subgroups with topranking probability of *Lactobacillus* and *Akkermansia* emerge. Finally, we plotted Quetelet index by subgroups to infer associations between subgroups and the cohort groups (Figure 6b). Quetelet index estimates the relative change of the occurence frequency of a latent subgroup among all the samples compared to that among the samples of the cohort groups. The index showed subgroups 1, 8,9, 10 are positively associated with AD group. The subgroup 9 is enriched by the members of *Ruminococcaceae* family. The top ranking *Ruminococcaceae_UCG_002* and *Akkermansia* are more abundant in AD group than the control group according to limma-voom analysis. *Akkermansia* overabundance in AD gut microbiota is counterintutive but was previously reported by others [25] and this genus is more abundant in the gut microbiota of Parkinson’s patients, also [49]. The subgroup 10, where *Escherichia/Shigella* is the top ranking genus, is strongly associated with AD group but negatively associated with other groups. Conversely, subgroups 2,4, and 7, which are enriched by short chain fatty acid producers, are positively associated with the control and MCI groups but negatively associated with AD.

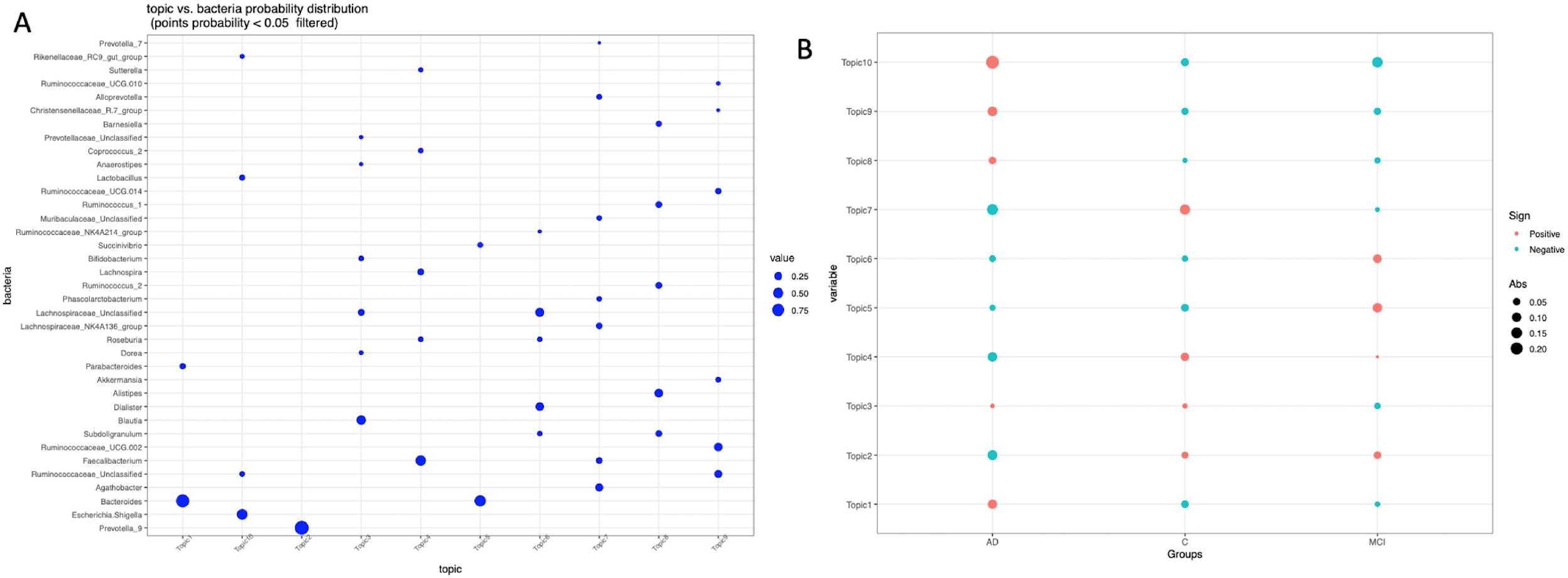

Another and last method we employed to stratify gut microbiota was topological data analysis (TDA), based on the *Mapper* algorithm [50] embedded in recently developed *tmap* tool [43]. The tmap tool was developed for network representation for stratification and association study of high-dimensional microbiome data. After constructing TDA microbiome network using Mapper algorithm (ordination, covering, and DBSCAN clustering) the workflow in the second step includes computation of a modified version of the spatial analysis of functional enrichment (SAFE) scores to map both the metadata and microbiome features into the TDA network to generate their vectors of SAFE scores. Vectors of SAFE scores are then used to perform ranking and ordination, and co-enrichment relations to delineate relationship between metadata and microbiome features. To construct TDA network we first applied dimension reduction (filtering) in PCoA using Bray-Curtis distance, followed the above algorithm and also repeated the entire analysis using Jensen-Shannon distance to reveal effect of distance metric, if any. To understand how driver taxa relate to each other and with the clinical metadata we performed Principal Component Analysis (PCA) of SAFE scores. Figure (7a) shows the TDA network and PCA (Bray-Curtis distance) of taxa-metadata based on SAFE scores (Supplementary Table S9), respectively. We obtained similar TDA network profile using Jensen-Shannon distance (Figures 7b) and SAFE scores Supplementary Table S10). Size of each marker is scaled according to the SAFE score and only top30 bacteria species are shown in PCA figures for clarity. A node in the network represents a group of samples sharing similar bacteria genus profiles. Two given nodes are linked when common samples are shared between the two nodes. The TDA analysis using both distance indices resulted in very similar stratification profile with the top ten SAFE scoring genera included *Prevotella_9, Bacteroides, Rumunococaceae_unclassified, species of Lachnospiraceae, and GCA90006675*. Unsurprisingly, a few taxa ranking differed between the two profiles such as *Caprococcus_2, Mollicutes_RF39_unclassified*.

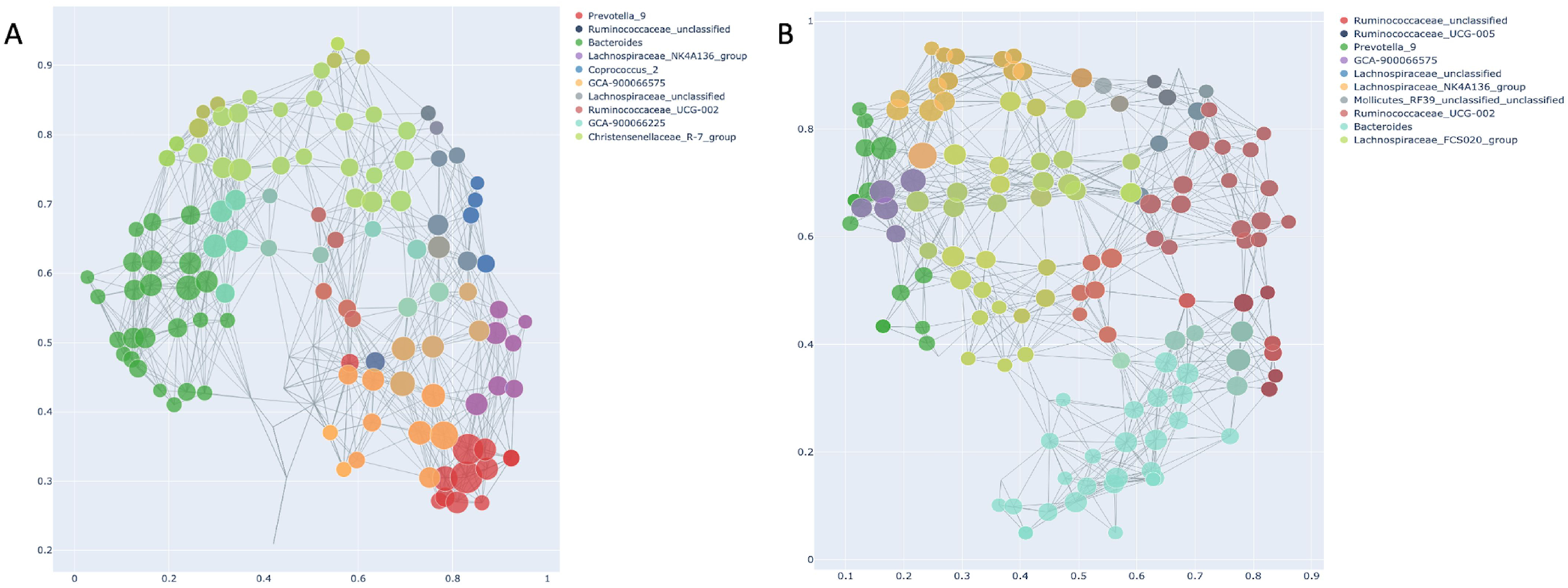

Furthermore, Figures (8a and 8b) show taxa and host covariates based on Bray-Curtis and Jensen-Shannon distances, respectively. Regardless of the distance metric, all three groups were clearly separated. The drivers of microbiome stratification (*Prevotella_9, Bacteroides, Ruminococcus_unclassified*) are placed near the control, AD and MCI groups, respectively in both PCA figures. Of the clinical metadata, MMSE, sex, and education were grouped with the control group and co-enriched with *Prevotella_9* but also with *Prevotella_2*, and *Haemophilus*, and *Lachnospiraceae_NK4B4_group*. Conversely, CDR, age, and AD group were clustered together and co-enriched with taxa such as *Subdoligranulum, Odoribacter, Bilophila, Alistipes*. The MCI group was co-enriched with *Ruminocoaceae_unclassified, Mollicutes_RF39_unclassified, Ruminocoaceae_UCG_005, Lachnospiraceae_unclassified*. However, some taxa such as *Odoribacter* was placed near the control group in Jensen-Shannon distance PCA Figure (8b), suggesting co-enrichment of certain taxa can be somewhat influenced by the preferred distance metric.

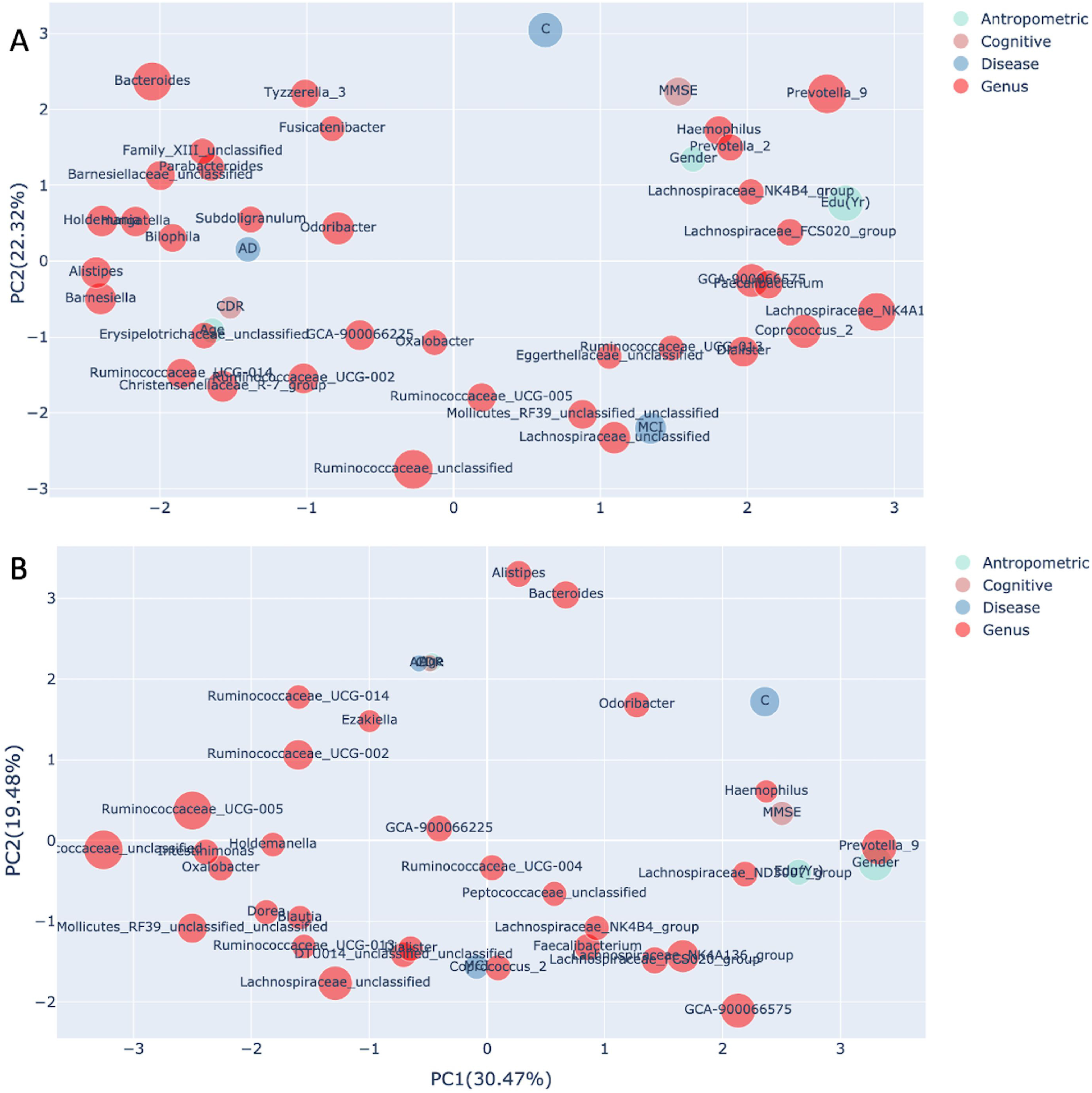

### Identification of signature taxa for AD continuum and association with metadata

We constructed Random Forest (RF) model on selected features of gut microbiota and psychometric test scores (MMSE and CDR) that are typically used as proxy in clinical diagnosis. Using songbird, we selected 300 ASV (Top 25%) that differentiates between the healthy (control) and the disease groups (MCI and AD). We then plotted the ASVs with the first 20 highest mean decrease Gini values (Figure 9a) and included ASVs with mean decrease Gini values above the breakpoint curve in the RF analysis. We identified the following 9 ASVs above the breakpoint: *Faecalibacterium (ASV45), Sutterella(ASV607), Coprobacter(ASV531), Bacteroides (ASV81), Anaerostipes(ASV364), Ruminoccocaceae_unclassified(ASV203), Lactobacillus (ASV65), Clostridium_sensu_stricto_1 (ASV118), Ruminococcus_1 (ASV59*). Notably, ASVs beyond the breakpoint line are largely the bacterial species responsible for the stratification of gut microbiota in the samples such as *Faecalibacterium, Bacteroides*, and *Ruminococcus_unclassified*. We next calculated diagnostic accuracy of the RF model by plotting receiver operating characteristics curve (ROC) for the above 9 taxa, MMSE, and CDR separately and in combination for each cohort group (Figure 9b). The ROC value for these selected nine taxa were moderately accurate (AUC 63%, FIg 8a) but when we included MMSE and/or CDR, we found that the RF model robustly classify all three groups (groupwise AUC range 0.74-1.0, Figures 9b).

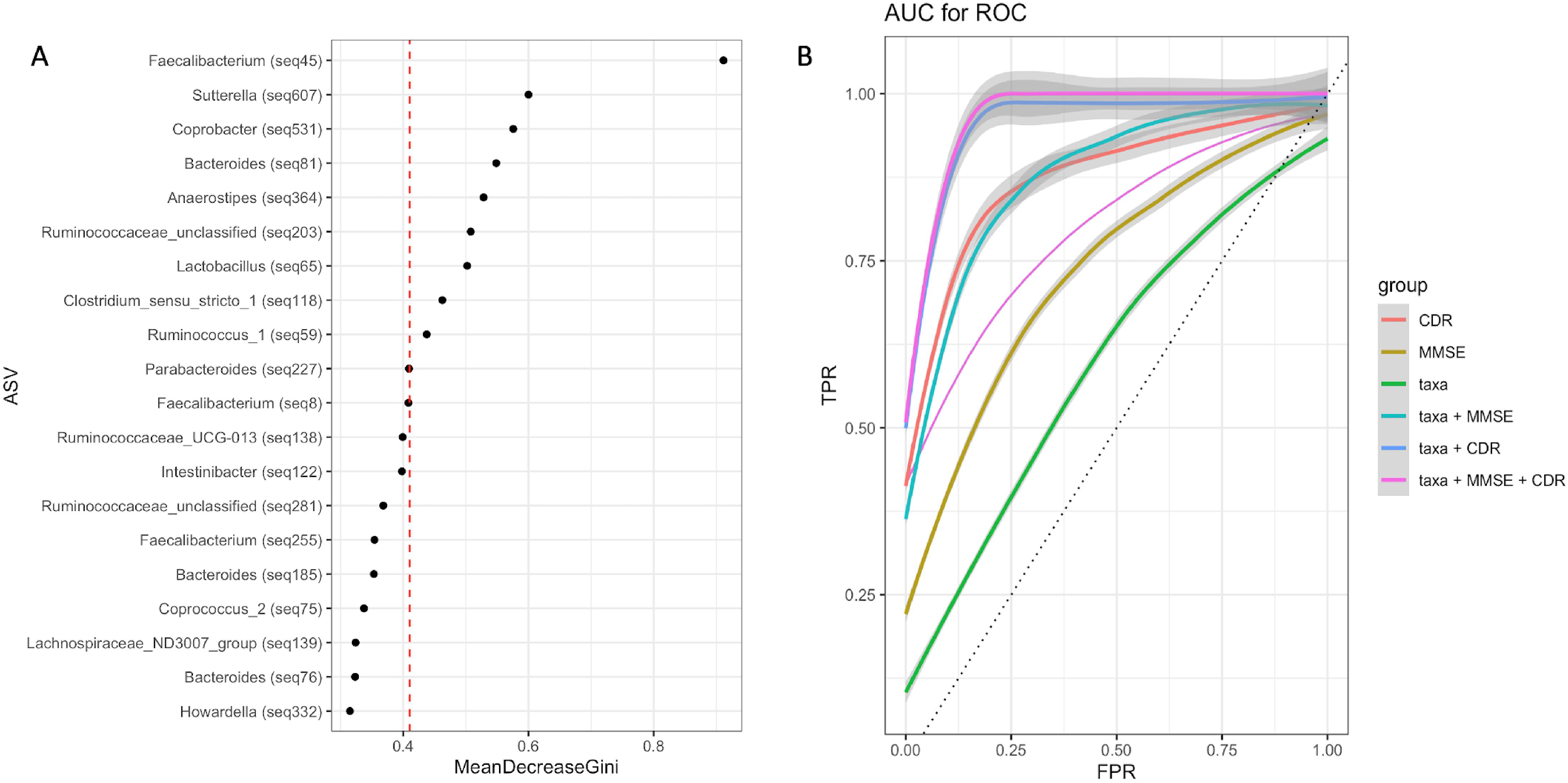

### Taxa association with clinical parameters

We used multivariate association with linear models (MaAsLin2) to assess association between individual taxa and clinical metadata including patients drugs (q ≤ 0.25). This analysis showed that *Roseburia, Lactobacillus, Fusicatenibacter* were negatively associated with AD (Supplementary Figure S5). Of the medication categories there are several taxa found to be positively associated with anti-depression and statin. *Blautia, Caprococcus, Butyricoccus, Dorea, Lachnospiraceae family members, some Ruminoclostridium and Ruminococaceae*, known to be butyrate producers are all positively associated with antidepression drugs. Unexpectedly, we found that several taxa were significantly associated with Statin medication and, of these taxa, *Streptococcus* and unclassified member of *Erysipelotrichaceae* were highly significantly associated with statin medication. We also observed the following taxa positively associated with statin medication; unclassified members of *Ruminococaceae* and *Lachnospiraceae, Phascolarctobacterium, Desulfovibrio, Caprobacter, Bifidobacterium, Butyricoccus, Blautia, Barnesiella*.

## DISCUSSION

In this study, we demonstrate that gut microbiota across AD continuum not only differentiates between cognitive states but also comprise subgroups delineated by locally dominant co-occurring bacteria. Stratification of the gut microbiota along the AD continuum is major unmet need for diet-based and precision nutrition interventions in AD cohorts and here we present proof-of-concept data that can be insighful for the emerging dietary and precision medicine/nutrition initiatives involving AD patients. A key finding in this study is that these approaches all converge on *Prevotella* and *Bacteroides* stratification, which are also robustly supported by enrichment and ordination analyses that these two species are the drivers of community diversity and composition. Rather than focusing on a single gut microbiota stratification method we have exercised the best practice of implementing multiple methods to compare, contrast, and sought support from alternative analyses. Also, all methods ranked the following taxa among the Top10 bacteria contributing to seperation of the groups; *Escherchia/Shigella, Faecalibacterium, Blautia, Ruminococcaceae_unclassified, Ruminococcaceae_UCG-002, Lachnospiraceae_unclassified, Parabacteroides*, suggesting these taxa play significant role in the observed community structure of the gut microbiota of the patients in this study.

PAM clustering and DMM concordantly showed three distinct clusters, one of which is consistent with the recently described Bact2 group [44]. The subjects in this group are likely to have aggravated dysbiosis as manifested from increased abundance of opportunistic pathogens *Escherichia/Shigella* and some species of *Bacteroides* species and lower abundance of *Faecalibacterium* and other SCFA producers. Notably, LDA analysis shuffles similar set of taxa as the number of subgroups increase but *Bacteroides* and *Prevotella_9* are predominantly the most abundant taxa in many of these clusters. Strikingly, *Escherichia/Shigella* dominates one of the subgroups in LDA analysis together with opportunistic *Klebsiella* and *Enterococcus*, suggesting dysbiotic community type may be enriched in this subgroup.

Topological data analysis (TDA) we used to stratify gut microbiota in this study deserves a particular attention among others. TDA, based on the Mapper algorithm [50], represents the underlying distribution of data in a metric space by dividing the data into overlapping similar subsets according to a filter function, local clustering on each subset and representing the results in an undirected network. A node in the network represents a group of samples with similar microbiome profiles, and if common samples between nodes are shared then the nodes are linked. Next, a modified special analysis of functional enrichment (SAFE) algorithm maps the metadata and taxa into the network. Finally, vectors of SAFE scores can be used in ordination to rank the driver taxa and their relationship with the metadata, all these algorithms are integrated into *tmap* [43]. The SAFE scores we obtain following these algorithms allowed us to identify the driver species that are responsible for community structure and showed their relationship with the metadata. We employed Bray-Curtis and Jensen-Shannon to check the variation resulting from distance metric. *Prevotella_9, Bacteroides*, and *Ruminoccus_unclassified* were ranked among the top10 taxa with high SAFE scores, albeit in different order, suggesting TDA is robust and consistent even with different distance metrics. In addition to these three taxa unclassified members of again other taxa within *Ruminoccus* family and *Lachnospiraceae* were congruent with other three methods we tested. Interestingly, this analysis identified *GCA-900066575* taxa (Uncultured human intestinal bacterium) as one of the subclusters in contrast with other methods we used. This genus is taxonomically in the family of *Lachnospiraceae*, which includes members of SCFA producers [51], still some other members were associated with metabolic diseases such as obesity [52]. Indeed, another related member of this family *GCA-900066225* ranked among the top10 taxa when Bray-Curtis distance was used but enriched around AD. It is therefore important to note that TDA, unlike clustering or probabilistic partitioning methods, provided fine resolution in terms of stratification of the gut microbiota composition. Conversely, TDA did not rank *Escherchia/Shigella* subnetwork among top ten taxa, neither the ordination showed clear association with the disease. Together, bioinformatic tools developed in the field of microbiome have all their strengths and drawbacks and therefore overlaps in bioinformatic analyses should be pursued.

Several lines of evidence showed human cohorts in microbiome studies can be phenotypically partitioned along *Prevotella* and *Bacteroides* stratification [53–58]. A recent comprehensive report [59] provided evidence that Mediterranean diet-based intervention is associated with specific functional and taxonomic components of the gut microbiome, and its effect is a function of microbial composition. Notably, absence of *Prevotella copri* in the gut microbiomes of a subgroup of participants was associated with the protective health benefits of the dietary intervention, emphasizing the premise that microbiome-informed stratified dietary intervention would be quite effective. Nevertheless, *P. copri* is ambivalently associated with both heath and diseases depending on the strain and geography [60], which prompts us to further consider its role in AD.

Taxonomically, the genus *Prevotella_9* is predicted to belong to *Prevotella copri* complex [61]. Comparative genome analysis of the strains of *P.copri* complex, however, show that some strains qualify to be assigned to even a separate species of *Prevotella* due to low genomic similarities [62, 63]. Some *P. copri* strains are associated with disease states such as rheumatoid arthritis [64], while some other strains are associated with habitual diet and life style [54] and underrepresented in Westernized populations. Thus, strain level resolution of *Prevotella_9* is needed to draw inferences. Expectedly, multiple strains of *P. copri* are likely to be part of the bacterial community in the samples. Even though we found *Prevotella_9* to be associated with the control group the enrichment analysis using songbird ranked some ASVs belong to Prevotella_9 (species level) at the top and few other ASVs at the bottom of the log ratio differentials, suggesting analysis beyond species taxonomic hierarchy would provide better resolution in terms of their associations with human phenotypes. Oligotypes of these two genera in an earlier work were found to be differentially associated with plant based or some others were associated with animal-based diet [55]. A recent report provided evidence that *Bacteroides cellulosilyticus* predicted weight gain more precisely than the ratio of *Prevotella* and *Bacteroides* genus. Together, our differential enrichment analysis results are in line with these reports that species or even strain level resolution of these two genera could provide better predictive biomarker power for diet-based intervention studies.

One limitation of our study was that although we were able control drug induced confounding, we did not control other potential confounders such as diet, BMI, stool consistency. We largely recruited cohabiting spouses as non-demented controls sharing the same diet patterns with the patients and carnivory is rare due to the high cost of meat in the country. We therefore did not predict diet can strongly impact our results.

In conclusion, we demonstrate in this study that gut microbiota along the Alzheimer’s Disease continuum comprises stratified community structure dashed primarily by *Prevotella* and *Bacteroides* but also subnetworks of other taxa exist in the community. The signature taxa when used together with MMSE and CDR robustly classify heterogenous groups hence posing potential biomarker value. The study adds to limited number of clinical studies profiling gut microbiota of AD continuum patients.

## MATERIALS AND METHODS

### Subject Recruitment and Study Design

The Istanbul Medipol University and Erciyes University Ethical Review Boards approved this study (Approval numbers: 186/16.4.2015 and 85/ 20.02.2015, respectively). All participants were informed of the objectives of this study and signed a written consent form prior to their participation. The diagnosis of dementia and MCI due to AD were based on the criteria of the National Institute on Aging-Alzheimer’s Association workgroups on diagnostic guidelines for Alzheimer’s disease [65, 66]. Exclusion criteria for this study included history of substance abuse, any significant neurologic disease, major psychiatric disorders including major depression. Also, individuals who used commercial probiotics or antibiotics during the study period or within 1-month prior to providing stool sample, or who major GI tract surgery in past 5 years. Both health centers followed the same protocols in recruiting cohorts and used kits from the same manufacturers to minimize the variations in wet lab procedures.

### Lumbar puncture, CSF biomarkers assays

Cerebro Spinal Fluid (CSF) samples were included in the analyses from a subset of AD patients if the patient was requested to donate CSF sample as part of the clinically mendated diagnostic protocol. CSF samples were collected in the morning after overnight fasting using spinal needles (22 gauge) and syringes at the L3/4 or L4/5 interspace. CSF was then aliquoted into 0.5 mL non-adsorbing polypropylene tubes and stored at −80 °C until assay. Biomarker molecules in CSF (Aβ_42_, phosphorylated tau (p-tau), and the p-tau/Aβ42 ratio) were measured consistent with the Alzheimer’s Association flowchart for lumbar puncture and CSF sample processing and the biomarker levels were determined as previously described [67]. Single 96-well ELISA kits (Innogenetics, Ghent, Belgium) were used in quantitation.

### *Sample collection* and *DNA* extraction

Stool samples from all participants were collected in the neurology clinics of the university training hospitals. The participants were given a collection kit included a sterile tube and provided a brief instruction for collection. Self-collected samples were placed within approximately 30 mins of collection in −80 freezers and kept frozen until DNA extraction.

16S rRNA gene sequencing and PCR were performed as previously described [68] with minor modifications. Briefly, genomic DNA was extracted from 220 mg fecal samples using QiaAmp DNA Stool Mini Kit (Qiagen, Germany) per manufacturer’s instructions with the addition of bead beating (0.1 mm zirconium-beads) and lysozyme and RNAse A incubation steps.

### PCR and amplicon sequencing

To amplify the variable V3-V4 regions of the 16S rRNA gene, the primers 341 F (5’-CCTACGGGNGGCWGCAG-3’) and 805 R (5’-GACTACHVGGGTATCTAATCC-3’) were used. MiSeq sequencing adaptor sequences were added to the 5’ ends of forward and reverse primers. Approximately 12.5 ng of purified DNA from each sample was used as a template for PCR amplification in 25 μl reaction mixture by using 2 × KAPA HiFi Hot Start Ready Mix (Kapa Biosystems, MA, USA). For PCR amplification, the following conditions were followed: denaturation at 95 °C for 3 min., followed by 25 cycles of denaturation at 95 °C for 30 sec., annealing at 55 °C for 30 sec. and extension at 72 °C for 30 sec., with a final extension at 72 °C for 5 min. Amplified PCR products were purified with Agencourt AMPure XP purification system (Beckman Coulter) and Nextera PCR was performed by using sample-specific barcodes. The constructed Nextera libraries were then sequenced by Illumina MiSeq platform using MiSeq Reagent Kit v2 chemistry.

### Sequence processing and taxonomic assignment

The pair-end 16S rRNA reads were first used cutadapt v1.9 program [69] for the process of quality filtering, trimming and uploaded on the DADA2 pipeline [34] integrated into the Nephele platform [70] (v.2.0, http://nephele.niaid.nih.gov). Chimeric sequences are automatically removed by this pipeline, which generates both rarefied and unrarefied ASV abundance tables. We used Rarefied (10769 reads/sample) ASV table in most downstream analysis due to large differences between some total sample reads except for the scale invariant DEICODE and songbird. We removed any sequences that were classified as either being originated from eukarya, archaea, mitochondria, chloroplasts or unknown kingdoms.

### Quality control

We included no sample DNA extractions and no template negative control samples in every sequencing library prepared. Using reads in the negative control samples as reference we identified and removed probable contaminant reads of 13 ASVs from the ASV table, as predicted by Decontam R package [71] using the ‘prevalence’ method. In this method, the binary coded features across samples are compared to the prevalence in negative controls to identify contaminants. Also, we sequenced the same amplicon of an AD sample three times to check the sequencing variation. Although both centers used same protocols and kits from the same manufacturer in sequencing, we sequenced amplicons amplified from two same genomic DNA templates again from AD samples at both centers to check the center-to-center sequencing concordance. No differences could be identified between the taxonomic compositions of the samples seuquenced at both centers nor between the technical replicates (PCoA, PERMANOVA p=0.1).

### Numerical Ecology and Statistical Analysis

Most numerical downstream analysis of ASV abundances were performed in R environment [72]. All P values, where appropriate, were adjusted for multiple testing using Benjamini-Hochberg (False Discovery Rate; FDR) method. We measured within samples microbial diversity (alpha diversity) using Observed richness, Chao1, Shannon, and Inverse Simpson in *phyloseq* [73] and tested using Kruskal Wallis. To identify differentially abundant bacterial species we employed animalculus [58] and limma [74] R packages. We assessed microbial diversity between samples (beta diversity) using multiple distance metrics including Bray-Curtis, Jaccard, Canonical Analysis of Principal Components (CAP). CAP analysis and the similarity percentages breakdown (SIMPER) procedure were performed using PRIMER.v7 [75]. Additionally, due to the compositional nature of the data, we also included robust Aitchison PCA, using the Qiime2 DEICODE plugin [36] to calculate beta diversity with feature loadings. The resulting ordination was visualized using Emperor [76]. We tested significance of beta diversity among groups using again Qiime diversity plugin PERMANOVA.

Next, we used Songbird [37] for multinomial regression to rank species association with disease status with the following parameters: (formula: “MMSE+CDR+Sex+Edu+C(Group, Diff, levels=(‘C’,‘MCI’,‘AD’), -p-epochs 10000 --p-differential-prior 0.5 --p-summary-interval 1 --p-random-seed 3 −min-sample-count 1000 −min-feature-count 0). Of note, the formula structure follows Patsy formatting (https://patsy.readthedocs.io/en/latest/) such that Groups (C, MCI, AD) represent levels=[“healthy”, “mild”, “severe”] states, respectively. A null model was generated using the same parameters. The fitted model demonstrated better fit compared to the null model (pseudo Q^2^ = 0.874027). Taxa ranks were visualized using Qurro [38]. Significance was determined using a Welch’s t-test between groups, performed by Graph Pad Prism.

To identify microbial species associated with the clinical metadata including patients’ medication we performed multivariate association with linear models (MaAsLin2) [77]. The control group was excluded from this analysis as they were not normally prescribed these medications. We employed the R package MaAsLin 2.1.0 to perform per-feature tests. We log-transformed relative abundances of microbial species and standardized continuous variables into Z-scores and binary encoded medication information before including them in the MaAsLin models (q<0.25 for significance).

### Stratification of gut microbiota

We employed clustering, probabilistic partitioning, and topological data analysis approaches for the stratification of gut microbiota in the samples. Partitioning around the medoid (PAM) approach [39] clusters samples by iteratively updating each cluster’s medoid. We assigned samples to community types using the function *pam(*) in R package *cluster* based on Bray Curtis and Jensen Shannon distances. The number of clusters was determined by Gap statistic evaluation. Departing from the clustering approach, we next used two distinct probabilistic methods to partition microbiota landscape, namely Dirichlet multinomial mixture models (DMM) [40] and Latent Dirichlet Allocation (LDA) [41, 42]. Genus level abundances were fitted to DMM models to partition microbial community profiles into a finite number of clusters, using the Laplace approximation as previously described [40, 78].

As a second probabilistic partitioning we performed LDA, is a multi-level hierarchical Bayesian model [41] otherwise used for collections of discrete data such as text corpus analysis in linguistics. LDA is a generalization of Dirichlet multinomial mixture modeling where biological samples are allowed to have fractional membership and distinct microbial communities have different microbial signatures. Thus, for each taxon there is a vector of probabilities across all clusters that it can be assigned to. Each cluster, therefore, has a different probability of containing taxa, indicating chance of microbes in a particular subgroup (strata) co-occurring due to community assembly dynamics. To fit the model we used Gibb’s sampling with the R package *MetaTopics* (v.1.0) [79]. The relative abundances of genus collapsed table with abundances more than 0.1% and 5% sample prevalence was input to the model. We plotted perplexity measure and loglikelyhood values to estimate model performance and optimal number of topics (subgroups of microbial assemblages) using 5-fold cross-validation. However, we observed that both parameters continued to improve with increasing subgroup number without a clear optimum except the first jump in perplexity was near 10 topics. We therefore picked first 10 topics for the sake of interpretability.

The final method we applied was topological data analysis (TDA) based on the Mapper algorithm [50] and network representation for stratification and association of study of high dimensional microbiome data, all integrated into *tmap* tool [43]. The framework enables to reveal association of taxa or metadata within the entire network and to identify enrichment subnetworks of different association patterns. Conceptually, the Mapper algorithm transforms a distance matrix and represent the shape of the data cloud in an undirected network. Next, a modified version of special analysis functional enrichment (SAFE) algorithm to map the value of the target feature into the network was employed, followed by ordination of SAFE scores to show taxa-metadata association [43].

### Signature taxa

To identify microbial signature of severity of cognitive impairment in AD continuum we implemented a machine learning procedure. We first took advantage of songbird tool to select features including the covariates and healthy (control) and disease states (AD+MCI) in the model formula. We subsequently fit the list of ASV selected this way into Random Forest models. We plotted the area under the receiver operating characteristic curve (AUROC) to show prediction performance of the models. To create the classifiers, a random forest constituted of 500 trees were computed using the default settings of the “randomForest” function implemented in the randomForest R package (v4.6-7). Mean decrease Gini values were averaged for each ASV among the 100 random forest replicates. The ASVs with the first 20 highest mean decrease Gini values were plotted. ASVs with mean decrease Gini values above the breakpoint curve were chosen to be part of the classifier. Breakpoints were estimated using the “breakpoints” function included in the strucchange R package [52]. We subsequently fit the list of ASVs selected this way with or without psychometric test values, i.e. MMSE and CDR, into Random Forest models, and bootstrapped for 100 times. We plotted the area under the receiver operating characteristic curve (AUROC) to show prediction performance of the models.

## DATA ACCESSION

The 16S rRNA generated by this study have been submitted to the NCBI BioProject database, (https://www.ncbi.nlm.nih.gov/bioproject/) under accession number PRJNA734525.

## AUTHOR CONTRIBUTIONS

**Conception and Design**: SY, OUN, EK, and LH; **Sample Collection and Processing**: BS, AG, FK, MFG, DK, AES, HAV, EAG, and KS; **Data Analysis**: SY, OUN, AB, MA, and MK; **Data Interpretation**: SY, OUN, MK, AM, LH and EK. **Manuscript Writing** – Original Draft: SY; **Writing, Review, and Editing**: OUN, MK, AM, LH and EK. All authors read and approved the final manuscript.

## LEGENDS FOR SUPPLEMENTAL TABLES AND FIGURES

**Table S1**. Levels of Cerebro-Spinal Fluid Biomarkers of a Subset of AD Patients.

**Table S2-S5**. Differentially Abundant ASV and genus level taxa between cohort groups as detected by Limma-Voom Model (Age and Sex Adjusted)

**Table S6**. PERMANOVA analysis of covariates

**Table S7-S8**. Enrichment analysis by multinomial regression embedded in the songbird (Set1, Set2, Set3, and Set4)

**Table S9-S10**: Ranking of SAFE scores calculated using tmap algorithm

**Figure S1**. **Alpha diversity analysis**. Box plots show (A) Chao1 index, (B) Inverse Simpson, (C) Observed species, (D) Shannon diversity index

**Figure S2**. **Multi-Dimensional Scale (MDS) Analysis of genus relative abundances**. (A) MDS analysis of the samples (B) Gradient of *Prevotella_9* abundances across the samples. (C) Gradient of *Bacteroides* abundances across the samples

**Figure S3. Determining the number of clusters in the gut microbiota.** The optimal number of clusters based on (A) Gap statistic with standart error bars for PAM analysis. (B) Laplace method for evaluating model fit for increasing number of Dirichlet mixture components

**Figure S4. Latent Dirichlet Allocation Model Performance.** LDA model’s perplexity parameter (top) and log-likelihood values (bottom) to find optimal number of clusters.

**Figure S5. Associations of the patient drugs with genus-level features.** The heatmap shows per-feature testing in MaAsLin 2 using linear mixed models to identify microbial species associated with drugs used by the patients. Colors of the heatmap reflects the beta coefficient for drugs and age and sex from linear mixed models in MaAsLin 2 with genus-level feature as outcomes.

## ACKNOWLEDGEMENTS

This study was supported by funding from The Scientific and Technological Research Council of Turkey (TÜBITAK) to Prof. Dr. Süleyman Yildirim and Prof. Dr. Emel Köseoglu (Project numbers. 2236-115C056 and 215S707, respectively). The funding agency had no role in study design, data collection and interpretation, or the decision to submit the work for publication.

